# Mice tails function in response to external and self-generated balance perturbation on the roll plane

**DOI:** 10.1101/2024.04.18.589832

**Authors:** Salvatore Andrea Lacava, Necmettin Isilak, Marylka Yoe Uusisaari

## Abstract

The functionality of mouse tails has been unexplored in the scientific literature, to the extent that they might seem to be considered as a passive appendage. Previous research on mouse locomotion has largely omitted tail dynamics, but hints at its potential use in balancing can be seen in the natural habitats and behaviors of these rodents. Here, leveraging high-speed videography, a novel naturalistic locomotory task and a simple biomechanical model analysis, we investigated the behavioral utility of the mouse tail.

We observed that mice engage their tails on narrow ridge environments that mimic tree branches with narrow footholds prone to roll-plane perturbations, using different control strategies under two defined conditions: during external perturbations of the ridge where they primarily work to maintain posture and avoid falling, and during non-perturbated locomotion on the ridge, where the challenge is to dynamically control the center of mass while progressing forward.

These results not only advance the existing understanding of mouse tail functionality but also open avenues for more nuanced explorations in neurobiology and biomechanics. Furthermore, we call for inclusions of tail dynamics for a holistic understanding of mammalian locomotor strategies.

**Author summary:** We describe and quantify the rapid mouse tail movements in response to external balance perturbations, possibly constituting a novel balance-compensatory motor program. Furthermore, we bring to light the consistent, context-dependent movements of the tail during increasingly precarious locomotion. The observations highlight the tail as an integral component of the mouse locomotory system, contributing to balancing and putatively movement efficacy, and call for inclusion of the tail in future works examining motor (dys)function.

## Introduction

Tails are a defining features of chordates, exhibiting significant diversity in both form and function [1]. For example, the prehensile tails of primates and rats have a range of utilities from grasping to signaling while the mouse tail seems a relatively little-used if not entirely passive appendage. It likely lacks the neuromechanical substrates for object manipulation and is rarely used in social signaling contexts, barring the occasional stress-induced ‘tail rattling’ [2]. The thin and weak mouse tail accounts for only a few percent of the body mass, seemingly negating its utility as a balancing counterweight during leaps, a function observed in lizards [3]. Its nearly hairless structure offers limited aerodynamic assistance in high-speed maneouvering in the manner of other light-tailed species such as cheetahs and squirrels [4]. Notably, studies analyzing body kinematics of mice during such high-speed locomotion and turns have predominantly excluded the tail from scrutiny, in line with a view of its limited role in locomotion [5, 6]. More recently the first modern 3D description of limb and tail kinematics of mouse locomotion on a broad, straight and stable surface [7] suggested that mice merely hold their tails out of their way without taking advantage of it.

Despite the seemingly limited functionality, the prevalent presence of tails in arboreal (squirrels, deer mice) but not in grassland rodents (prairie vole, hamster, prairie dog) points to a potential utility in balancing — a role that, surprisingly, has not been rigorously investigated. In fact, the “mouse balancing task” has primarily served as a proxy for assessing brain or body function, such as the severity of ataxia [8] or spinal cord injury [9], with virtually no studies examining the specific motor strategies mice employ to maintain balance. Only recently the neuromechanical substrate of balancing function has been brought to experimental scrutiny [9, 10], but the focus still has been on limb-rather than tail-based strategies. Strikingly, despite the wide use of balance beam crossing task [11] and availability of immense amounts of video recordings from such experiments, no systematic descriptions of mouse tail movements during the task have been provided.

This is intriguing given that mice (Mus musculus) naturally navigate complex environments and exhibit a level of arboreal behavior, and thus the tail would be expected play an active role in balance and stability in contexts with narrow footholds where the animal is particularly vulnerable to roll-plane (sideways) perturbations.

Leveraging high-speed videography and quantitative analysis, we provide the first unequivocal demonstration and quantification, to our knowledge, of the behavioral utility of mouse tails in balance maintenance. Our starting observation is familiar to all those who routinely handle mice: when held in experimenter’s hand and subjected to roll-plane rotations, mice often rotate their tails swiftly in a direction counter to the imposed movement. This tail-rotation response is also reproduced when a mouse is placed in a motorised, roll-plane swing (Figure 1a; S1 Video). These observations highlight the tail as an integral component of the mouse locomotory system, contributing to balancing and putatively movement efficacy, and call for inclusion of the tail in future works examining motor (dys)function.

**Figure 1.**
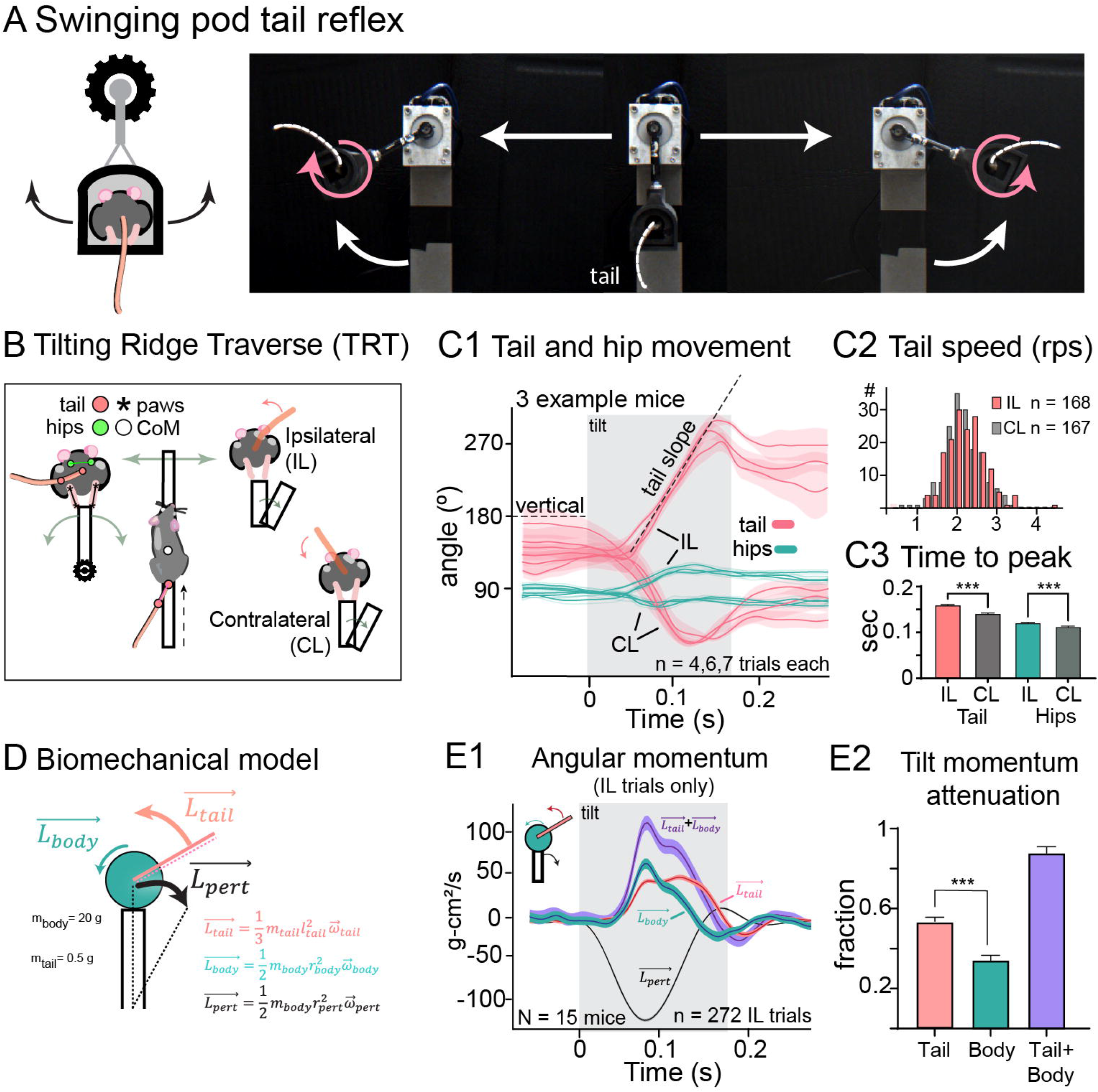
Roll-plane tilting evokes counteracting tail responses. A: mice experiencing lateral rolls in a custom-made swing rotate their tails in the countering direction. Schematic illustration (left) and screenshots from S1 Video (right). B, schematic illustration of the Tilting Ridge Traverse (TRT) task. Left: location of tracked body parts as seen from rear and top cameras. Right: depiction of “ipsilateral” and “contralateral” trials, defined by whether the tilt occurred in the same or opposite direction of the tail position, respectively. C, tail and hips angle changes in response to tilts. C1, tail and hip angle trajectories (pink and green data, respectively) from 3 example mice experiencing either ipsilateral (IL) or contralateral (CL) tilts. Traces show mean *±* SEM for n = 4, 6 and 7 trials. Gray area indicates ridge movement. C2, comparison of mean tail acceleration (defined as the slope as indicated in C1) in IL and CL trials. C3, comparison of tail and hip movement duration (to the maximal displacement) in IL and CL trials. D, schematic illustration of the biomechanical model used to estimate angular momenta experienced by the mouse as well as the compensation generated by body and tail rotation based on tracked movements. For details, see Methods. E, comparison of angular momenta generated by the tilt perturbations and tail and body rotations elicited by IL tilts. E1, instantaneous momenta estimated using the model. Traces show mean *±* SEM for n = 272 trials N = 15 mice for tail, body and the sum of tail and body, as well as the estimated rotational momentum generated by the perturbation. Downwards direction corresponds to the direction of the tilt. E2, total angular momentum generated by tail, hips and their sum, shown as a proportion of the perturbatioin-generated momentum. All data shown as mean ± SEM and groups were compared using t-test (d) or one-way ANOVA (e) followed by Bonferroni’s post-test (*** p *<* 0.001).

## Results

### Rotational movement of tail generates significant angular momentum

In order to describe the balancing strategies mice use in naturalistic behavior, we developed a novel experimental task (“Tilting ridge traverse”, TRT; Figure 1B; S2 Video). The TRT task aims to mimic conditions that mice may encounter in nature while traversing a thin branch that is suddenly perturbed. In the task, the ridge is tilted left or right during randomly-chosen trials to elicit balancing responses that are recorded with two high-speed video cameras positioned above and behind the animal.

While traversing the ridge, the mice generally hold their tails on one side of the ridge and we first examined TRT trials where the tilt occurred in the direction of tail (“Ipsilateral (IL) tilt”; Figure 1B), expecting to observe body posture adjustment that would shift mass in the opposite direction. Indeed, the tilt elicited not only a moderate back-and-forth motion of the hips (Figure 1C1, green data) but also a rapid tail swing to the opposite side with remarkably invariant kinematics (Figure 1C, red data).

To our surprise, tilts that were directed to the opposite side of the tail (“Contralateral (CL) tilt”; Figure 1B) also elicited tail responses with mirrored kinematics and nearly-identical speed profiles (IL, 2.20 ± 0.037 rotations per sec (rps); CL, 2.12 ± 0.038 rps; p = 0.15) even though the range of movement was limited by the tail making contact with the ridge (Figure 1C1-C3; see Table 1). As in CL tilt trials the tail positioning can not provide additional counterweight, the main mechanism by which the tail supports balancing could relate to rotational momentum rather than simple weight shifting.

**Table 1.**
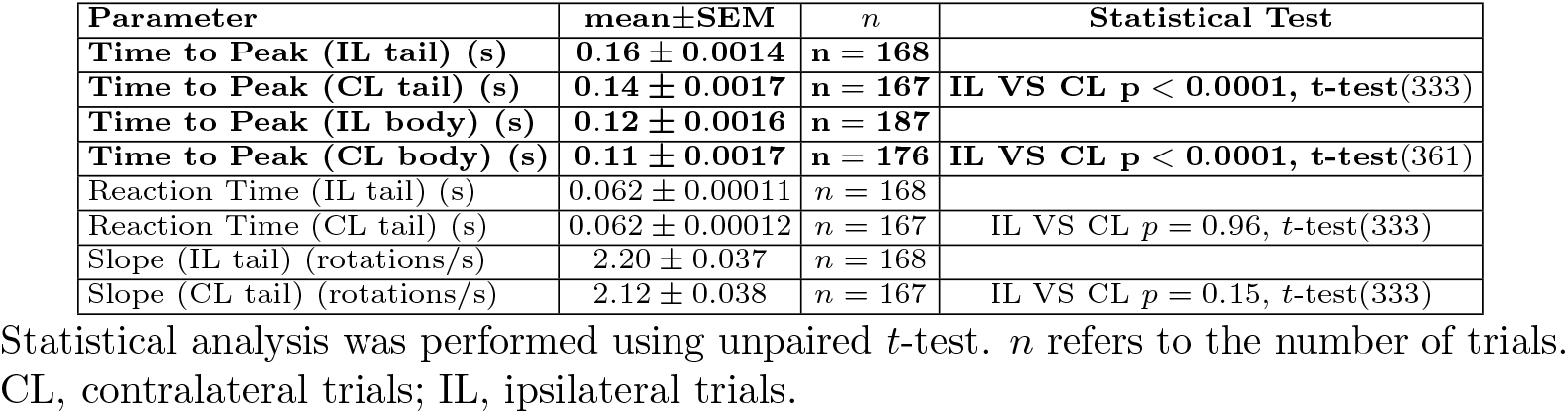
Tail kinematic measurements.

To investigate whether the tail angular momentum generated by the swinging motion could theoretically provide adequate compensation, we constructed a simple mechanical model that approximates the mouse body as a cylinder and the tail as a stiff rod attached to the base of the cylinder (Figure 1D; see Methods for model details). Using body part weights measured from the carcasses and angular velocities from experimental animals we estimated the angular momenta that our ridge-tilt conditions would instigate in the body as well as the theoretical compensation that the tail and body rotation could provide. As the contralateral tail movements were limited due to contact with the ridge (evidenced in the lower movement range in Figure 1C) and the resulting compensatory momentum greatly depends on the position of the tail at the onset of the ridge tilt, we only consider ipsilateral trials for the rest of this work.

Despite weighing only a fraction of the body mass (2.56 ± 0.12 % of the body without a tail), the observed movement of mouse tail results in compensatory momentum with peak magnitude comparable to that generated by rest of the body (Figure 1E1). Even more, the total tail-generated momentum was larger than that of the body during the tilt (Figure 1E2). This was due to the high speed of the tail (corresponding to up to 6 full rotations per second) as well as to the fact that the the tail rotation continues nearly throughout the 190 ms tilting motion while the hip movement terminates earlier (time to peak position for tail and hips during IL tilts 0.16 ± 0.0014 and 0.12 ± 0.0016 s, unpaired two-sample t-test *p <* 0.001). Ultimately, the sum of compensatory momentum by body and tail amounted to over 80 % of the total estimated rotational momentum experienced by the body in response to the ridge tilt (Figure 1E2, blue; see Table 2).

**Table 2.**
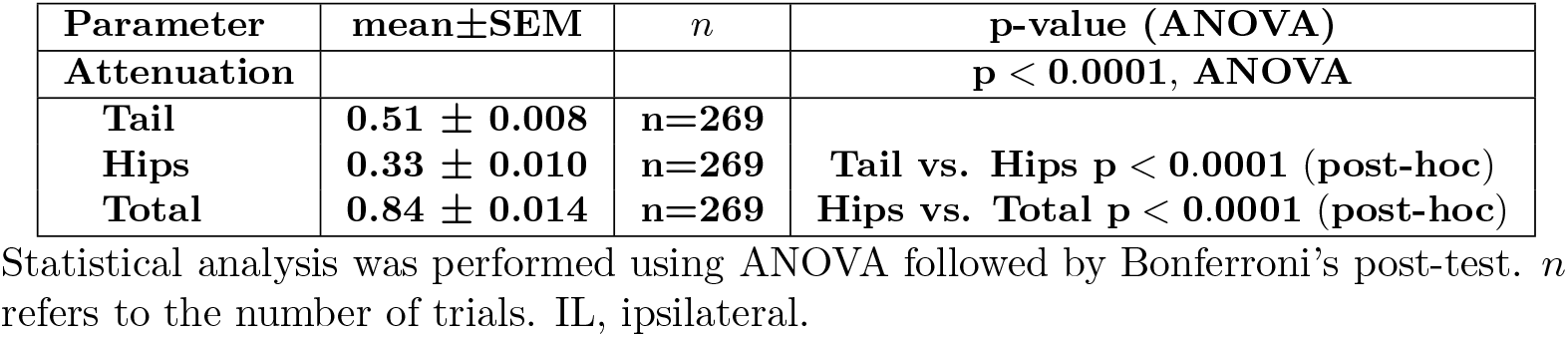
Angular Momentum Attenuation with Respect to Tilt (IL trials)

### Adjustments of the tail response in increasingly challenging conditions

Next, we investigated how the relative contributions of the tail and body are affected when the magnitude of perturbation (tilt angle) is either increased (to 30°; ‘L’ tilt) or decreased (to 10°; ‘S’ tilt; respective durations of the tilts 150, 190 and 230 ms; Figure 2A1; S3 Video). To provide a metric for the relative challenge posed by different tilts, we quantified the effects of the tilts on forward movement of the mouse (Figure 2A2-4). The performance of mice experiencing either 10 or 20-degree tilts were indistinguishable, but the largest (30 degree) tilt posed a greater challenge and lead to complete stopping (defined as forward speed less than 1mm/s, previously defined as immobility threshold in [12]) and a significant reduction in the distance the mice traversed in the 0.5 sec time window after the tilt (Figure 2A3-4; Table 3). Thus, mice were unable to completely compensate for the largest tilt perturbation and struggled to maintain forward movement. Indeed, when examining the time courses and magnitudes of tail and body-originating compensatory momenta, we found that even though the tail swing duration extended slightly with medium tilts compared to short tilts (Figure 2B1, red), with long tilts the tail movement was limited by contact with the opposite surface of the ridge. Thus, the total compensatory momenta could not increase further (Figure 2B2) and, in absence of modulation of the hip movements, led to incomplete compensation of the perturbation with long tilts (Figure 2B3).

**Table 3.**
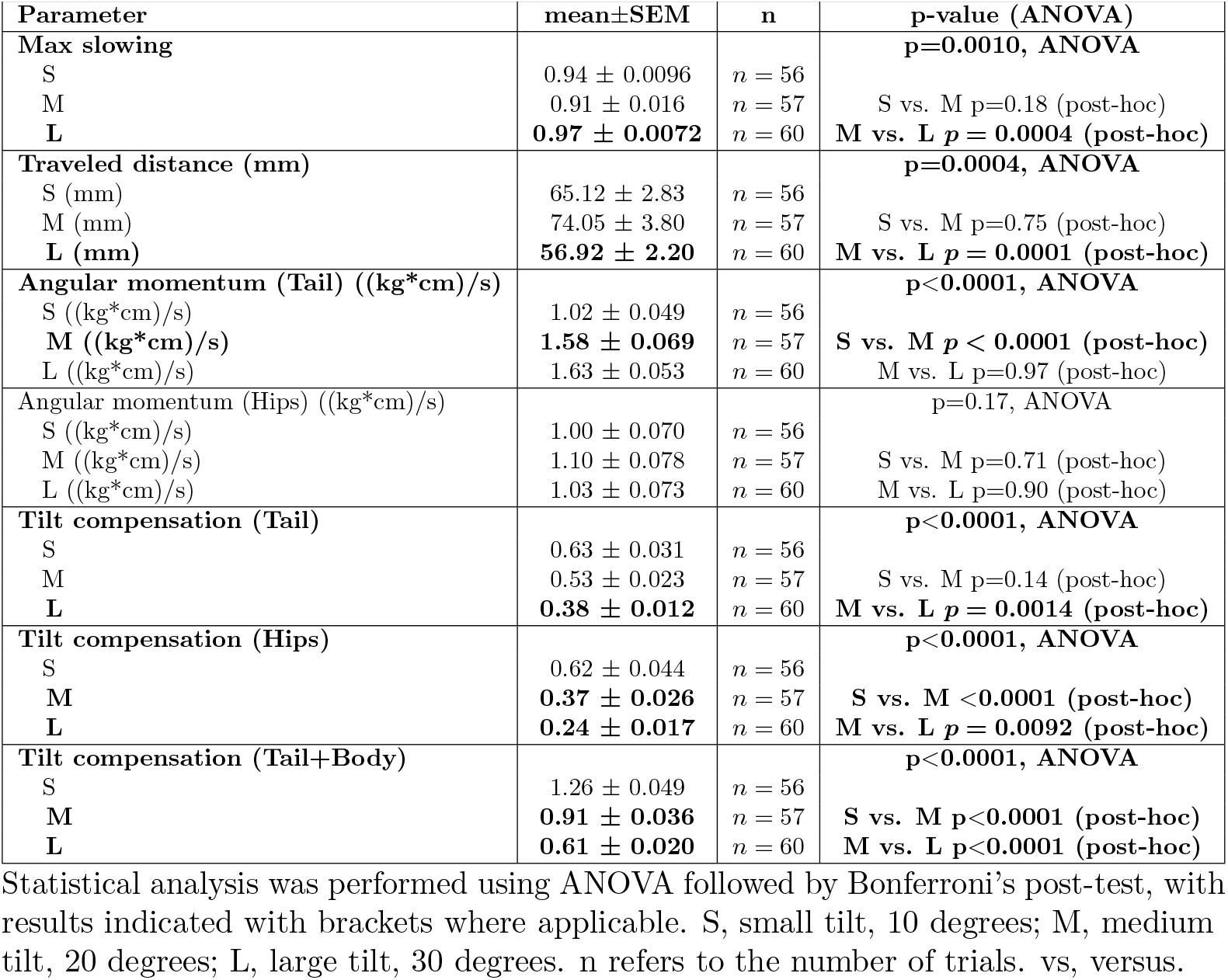
Effect of varying tilt amplitude.

**Figure 2.**
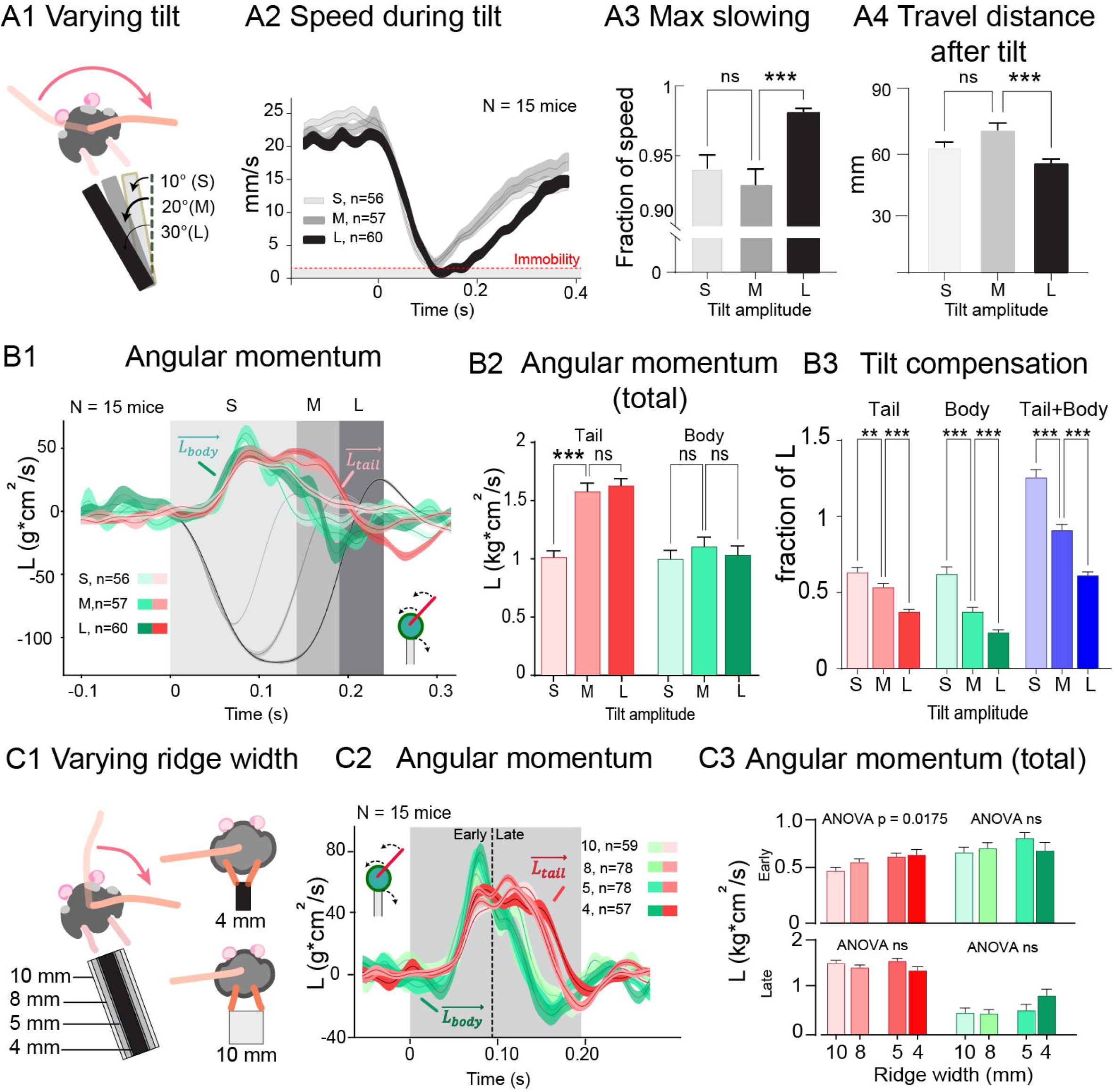
Tilt duration and ridge width effect on tail response to external perturbations. A, effect of tilt amplitude on balancing performance. A1, three tilt angles used: short (S), medium (M), long (L). A2, decrease in forward velocity of the mice during perturbations. Dashed line indicates threshold value for immobility (1 mm/s). A3, effect of larger tilt on task performance quantified as extent of slowing from pre-tilt-velocity. A4, distance travelled during the 0.5 sec following a tilt. Data from 15 mice; number of trials on each tilt amplitude indicated in panel A2. B, effect of tilt amplitudes on tail and body responses. B1, time course of the tail (red traces) and body-generated (green traces) angular momenta opposing the tilt-induced momenta (gray traves). Shading denotes tilt durations and the time windows from which total momenta are calculated in B2-B3. B2, total angular momenta generated by the tail and body during tilts. B3, total momenta for the tail, body, and their sum as a fraction of the total tilt-induced momentum. C, narrowing stance on ridge leads to slight changes in tail swing response. C1, schematic depicting different alignment of hind paws on narrow and wide ridges. C2, time course of tail and body angular momenta in response to medium-duration tilts on ridges of different widths. Shaded area indicates tilt duration; dashed line denotes division into early and late halfs used in C3. C3, total momentum of the tail response increase on narrower ridges during early phase of the response (top panels) but not late phase (bottom panels). Data are presented as the mean ± SEM, and statistical comparisons were conducted using one-way ANOVA followed by Bonferroni’s post-test (*p*<*0.05, **p*<*0.01, and ***p*<*0.001).

The kinematic invariance (i.e. invariance of speed profile) of the tilt-evoked tail responses to both ipsi- and contralateral tilts (as shown in Figure 1C) suggested that it might constitute of a previously-overlooked balancing reflex, analogous to other balance-related corrective motor programs [10, 13]. To investigate whether the tail swing and body rotation responses are affected by proprioceptive context, we repeated the experiments using ridges of different widths (4, 5, 8 and 10 mm) that force the mice to walk with subtly but significantly different postures (Figure 2c1; S4 Video). Focusing on the early phase of the compensatory responses that is most likely to be affected by the different body configuration, we found only a slight trend towards higher tail-generated compensatory momentum on narrow ridges (ANOVA *p <* 0.5); Figure 2C3; Table 5) during the first half of the response, while no effects were seen in momentum generated by body rotation. No differences were found either in body or tail-originating momenta in the late phase of the response.

**Table 4.** Effects of varying ridge width during tilted trials.

| Parameter | mean±SEM | n | p-value (ANOVA) |
| --- | --- | --- | --- |
| **Early phase ang mom (Tail) ((kg\*cm)/s)** |  |  | **p=0.0175, ANOVA** |
| 4mm ((kg*cm)/s) | 0.66 ± 0.050 | n = 57 |  |
| 5mm ((kg*cm)/s) | 0.65 ± 0.032 | n = 78 | 4 vs. 5 p=0.98 (post-hoc) |
| 8mm ((kg*cm)/s) | 0.59 ± 0.029 | n = 78 | 5 vs. 8 p=0.57 (post-hoc) |
| 10mm ((kg*cm)/s) | 0.51 ± 0.032 | n = 59 | 8 vs. 10 p=0.27 (post-hoc) |
| **Early phase ang mom (Body) ((kg\*cm)/s)** |  |  | **p=0.28, ANOVA** |
| 4mm ((kg*cm)/s) | 0.70 ± 0.077 | n = 57 |  |
| 5mm ((kg*cm)/s) | 0.82 ± 0.050 | n = 78 | 4 vs. 5 p=0.35 (post-hoc) |
| 8mm ((kg*cm)/s) | 0.73 ± 0.051 | n = 78 | 5 vs. 8 p=0.46 (post-hoc) |
| 10mm ((kg*cm)/s) | 0.69 ± 0.047 | n = 59 | 8 vs. 10 p=0.94 (post-hoc) |
| **Late phase ang mom (Tail) ((kg\*cm)/s)** |  |  | **p=0.077, ANOVA** |
| 4mm ((kg*cm)/s) | 0.68 ± 0.037 | n = 57 |  |
| 5mm ((kg*cm)/s) | 0.78 ± 0.026 | n = 78 | 4 vs. 5 p=0.051 (post-hoc) |
| 8mm ((kg*cm)/s) | 0.71 ± 0.024 | n = 78 | 5 vs. 8 p=0.22 (post-hoc) |
| 10mm ((kg*cm)/s) | 0.76 ± 0.030 | n = 59 | 8 vs. 10 p=0.63 (post-hoc) |
| **Late phase ang mom (Body) ((kg\*cm)/s)** |  |  | **p=0.78, ANOVA** |
| 4mm ((kg*cm)/s) | 0.41 ± 0.066 | n = 57 |  |
| 5mm ((kg*cm)/s) | 0.26 ± 0.062 | n = 78 | 4 vs. 5 p=0.15 (post-hoc) |
| 8mm ((kg*cm)/s) | 0.22 ± 0.040 | n = 78 | 5 vs. 8 p=0.94 (post-hoc) |
| 10mm ((kg*cm)/s) | 0.23 ± 0.050 | n = 59 | 8 vs. 10 p=0.99 (post-hoc) |

Statistical analysis was performed using ANOVA followed by Bonferroni's post-test. n refers to the number of trials. vs., versus.

**Table 5.**
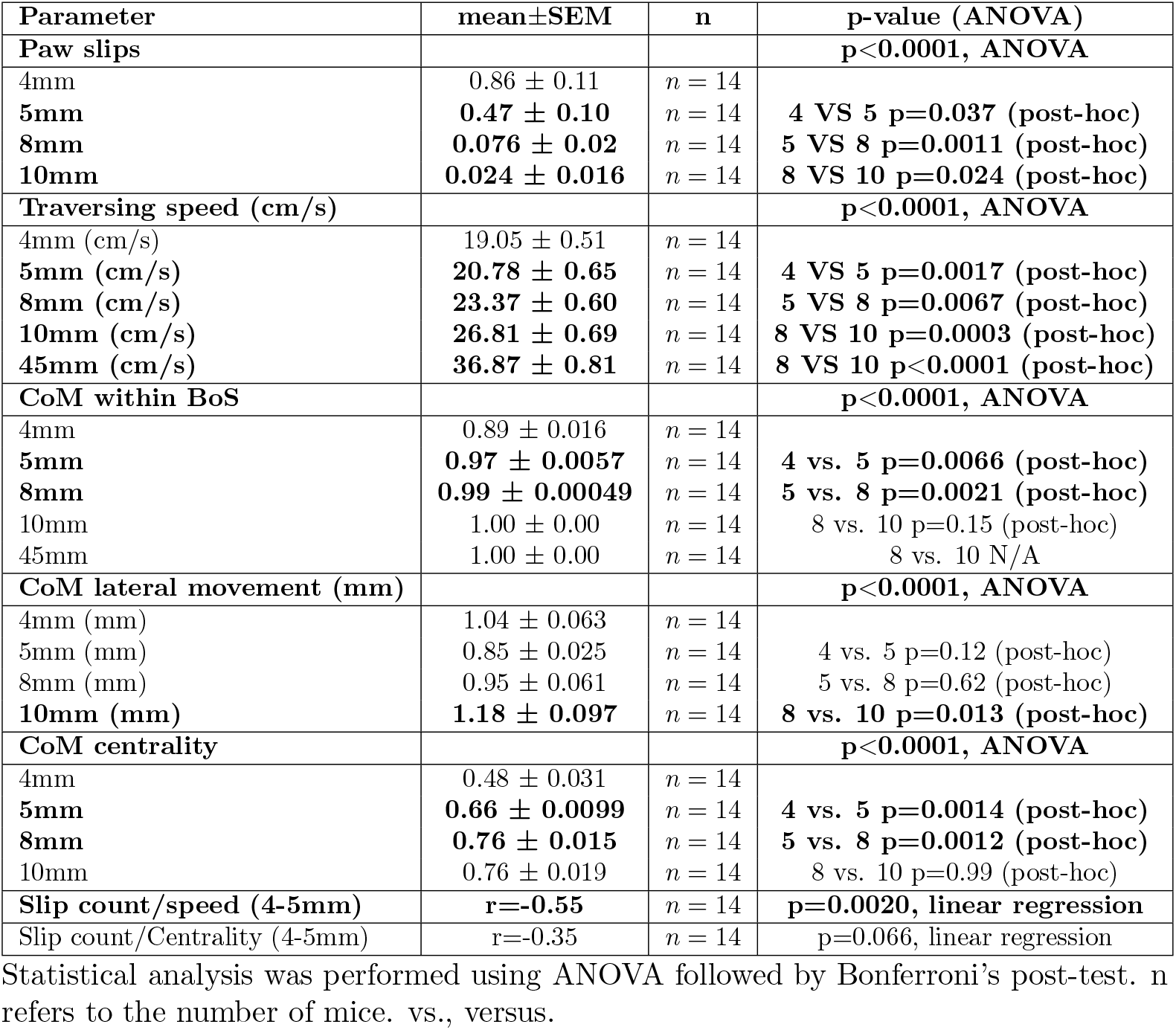
Locomotory performance on ridges with different widths.

| Parameter | mean±SEM | n | p-value (ANOVA) |
| --- | --- | --- | --- |
| **Paw slips** |  |  | **p<0.0001, ANOVA** |
| 4mm | 0.86 ± 0.11 | n = 14 | |
| 5mm | **0.47 ± 0.10** | n = 14 | **4 VS 5 p=0.037 (post-hoc)** |
| 8mm | **0.076 ± 0.02** | n = 14 | **5 VS 8 p=0.0011 (post-hoc)** |
| 10mm | **0.024 ± 0.016** | n = 14 | **8 VS 10 p=0.024 (post-hoc)** |
| **Traversing speed (cm/s)** |  |  | **p<0.0001, ANOVA** |
| 4mm (cm/s) | 19.05 ± 0.51 | n = 14 | |
| 5mm (cm/s) | **20.78 ± 0.65** | n = 14 | **4 VS 5 p=0.0017 (post-hoc)** |
| 8mm (cm/s) | **23.37 ± 0.60** | n = 14 | **5 VS 8 p=0.0067 (post-hoc)** |
| 10mm (cm/s) | **26.81 ± 0.69** | n = 14 | **8 VS 10 p=0.0003 (post-hoc)** |
| 45mm (cm/s) | **36.87 ± 0.81** | n = 14 | **8 VS 10 p<0.0001 (post-hoc)** |
| **CoM within BoS** |  |  | **p<0.0001, ANOVA** |
| 4mm | 0.89 ± 0.016 | n = 14 | |
| 5mm | **0.97 ± 0.0057** | n = 14 | **4 vs. 5 p=0.0066 (post-hoc)** |
| 8mm | **0.99 ± 0.00049** | n = 14 | **5 vs. 8 p=0.0021 (post-hoc)** |
| 10mm | 1.00 ± 0.00 | n = 14 | 8 vs. 10 p=0.15 (post-hoc) |
| 45mm | 1.00 ± 0.00 | n = 14 | 8 vs. 10 N/A |
| **CoM lateral movement (mm)** |  |  | **p<0.0001, ANOVA** |
| 4mm (mm) | 1.04 ± 0.063 | n = 14 | |
| 5mm (mm) | 0.85 ± 0.025 | n = 14 | 4 vs. 5 p=0.12 (post-hoc) |
| 8mm (mm) | 0.95 ± 0.061 | n = 14 | 5 vs. 8 p=0.62 (post-hoc) |
| 10mm (mm) | **1.18 ± 0.097** | n = 14 | **8 vs. 10 p=0.013 (post-hoc)** |
| **CoM centrality** |  |  | **p<0.0001, ANOVA** |
| 4mm | 0.48 ± 0.031 | n = 14 | |
| 5mm | **0.66 ± 0.0099** | n = 14 | **4 vs. 5 p=0.0014 (post-hoc)** |
| 8mm | **0.76 ± 0.015** | n = 14 | **5 vs. 8 p=0.0012 (post-hoc)** |
| 10mm | 0.76 ± 0.019 | n = 14 | 8 vs. 10 p=0.99 (post-hoc) |
| **Slip count/speed (4-5mm)** | **r=-0.55** | n = 14 | **p=0.0020, linear regression** |
| Slip count/Centrality (4-5mm) | r=-0.35 | n = 14 | p=0.066, linear regression |

Statistical analysis was performed using ANOVA followed by Bonferroni's post-test. n refers to the number of mice. vs., versus.

These observations suggest that the utility of the fast-swinging tail could increase in situations with strong perturbations (here presented as longer tilts or more demanding posture) where body rotation can not provide further compensatory rotation.

### Balance performance and mouse body posture

The differences in tail kinematics on thinnest ridges are likely reflecting increasing balancing challenges, exemplified by the gradual and significant increase in slips during traverse as well as a decrease in traversing speed in trials without tilt perturbation (Figure 3A1-2; S5 Video; Table 5). To gain deeper insight into how mice use their tails to support such precarious locomotion, we needed to construct finer metrics of balancing performance as a discrete count of paw slips might not be informative enough. Thus, we complemented the paw slip counts with additional metrics based on the center of mass (CoM) position with respect to the lateral extent of the base of support (BoS; see Methods and Table 5.

**Figure 3.**
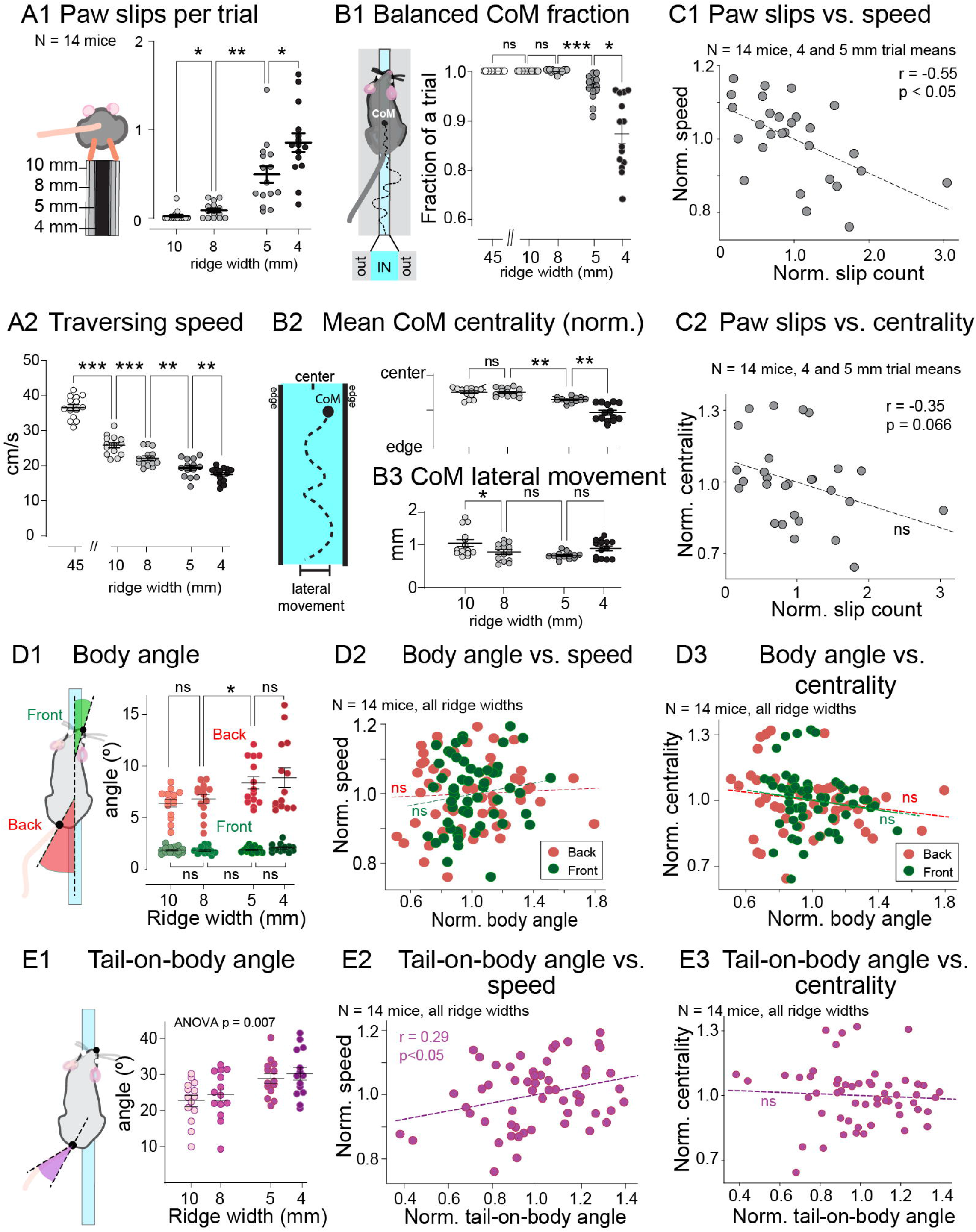
Effect of ridge width on body posture and task performance. A, classical quantification of ridge traverse performance by number of paw slips (A1) and traversing speed (A2). B, ridge traverse performance quantification based on Center of Mass position. B1, percentage of frames in which the Center of Mass (CoM) was entirely within the Base of Support (BoS), as determined by the ridge edges (schematic on the left). B2, left: schematic depiction of CoM centrality measure that ranges from 1 (at center of the ridge) to 0 (at or beyond the edge). Right, top panel: CoM position is more precarious (less central) on narrow ridges. Bottom panel: amplitude of lateral CoM movement does not differ on narrowest ridges. C, comparison of balancing performance metrics. C1, paw slip counts versus traversing speed; C2, paw slip counts versus CoM centrality. As very few paw slips occur on 8 and 10 mm ridges, only 5 and 4 mm trials are shown. D, effect of ridge widths on body alignment. D1: schematic (left) and summary of mean alignment angles of front and hind-body with respect to the ridge. D2: posture adjustment angle does not correlate with traversing speed. D3, increasing body angles (red, back; green, front) correlates with slightly less central CoM position. E, tail-on-body alignment on different ridge widths. E1, schematic (left) and summary of mean tail angles with respect to the hind-body angle. E2, larger tail-on-body angles were correlated with better performance in terms of traversing speed. E3, tail-on-body angles do not correlate with CoM centrality. In C, D and E, the values are normalized to ridge width-group means. All data are presented as mean ± SEM and were subjected to statistical analysis using one-way ANOVA followed by Bonferroni’s post-test (* *p <* 0.01, ** *p <* 0.001, and *** *p <* 0.0001).

As shown in Figure 3B2, the animals were able to maintain their CoMs above the ridge (acting as base of support, BoS) throughout nearly all and entire trials except on the most narrow (4 mm) ridge. On the 5 mm ridge the CoM stayed veered outside the support base very rarely, and even on the most narrow ridge the animals maintained balance during most of the trial (89 *±* 0.16% of the frames; see Table 5 for all values). A complementary and finer measure (“Relative centrality”; Figure 3B2) was based on the notion that for most energy-efficient locomotion the mouse should strive to maintain its CoM as close as possible to the center of the BoS. Again, we found that mice have no difficulties in maintaining their bodies close to midline on ridges broader than 5 mm. However, the centrality measure shows that on 5- and 4-mm ridges the CoM position becomes significantly more precarious, as the locomotion-related lateral oscillation of CoM can not be decrease further (Figure 3B3).

Importantly, the narrower footholds (see schematic in Figure 2C) on 4- and 5 mm ridges forced the mice to adjust hind paw posture. Furthermore, even though on all ridges the animals maintained the alignment of their head and front body with the ridge (Figure 3D1, green), on 5 and 4 mm ridges the caudal body posture became angled, (Figure 3D1, red; see also S5 Video), likely due to the challenge of placing paws on narrow support. However, the angles of either front or hind-body were not associated with slower advanging despite slight and nonsurprising correlation with CoM centrality (Figure 3D2-3).

In addition to the angled hind-body posture, mice also held their tails at larger angles with respect to the hind body when traversing 4-5 mm ridges (Figure 3E1). As the capacity for generating rotational momentum increases with the angle of the rotating mass [14], such adjustment could be part of a motor strategy to compensate for the increasing challenge of locomotion on narrower ridges. Indeed, mice traversed the ridges faster during trials in which they held their tails tail at high angles (normalized to ridge-group means; Figure Figure 3E2). However, as there was no effect of the tail angle on CoM centrality (Figure 3E3; see Table 6 for all values), the average tail position might contribute to biomechanical efficacy of locomotion rather than maintaining balance.

**Table 6.**
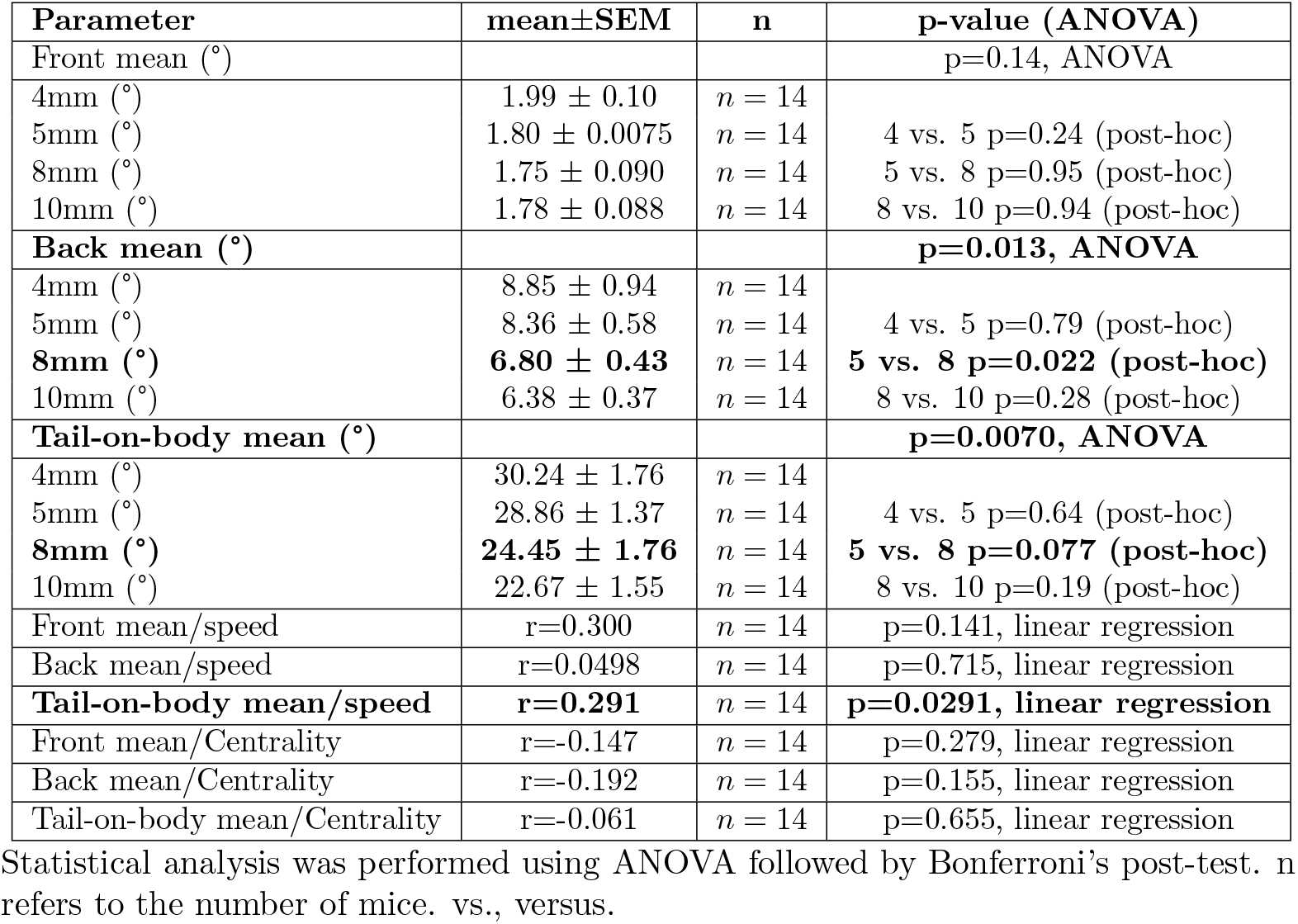
Posture on ridges with different widths.

| Parameter | mean±SEM | n | p-value (ANOVA) |
| --- | --- | --- | --- |
| Front mean (°) |  |  | p=0.14, ANOVA |
| 4mm (°) | 1.99 ± 0.10 | n = 14 | |
| 5mm (°) | 1.80 ± 0.0075 | n = 14 | 4 vs. 5 p=0.24 (post-hoc) |
| 8mm (°) | 1.75 ± 0.090 | n = 14 | 5 vs. 8 p=0.95 (post-hoc) |
| 10mm (°) | 1.78 ± 0.088 | n = 14 | 8 vs. 10 p=0.94 (post-hoc) |
| **Back mean (°)** |  |  | **p=0.013, ANOVA** |
| 4mm (°) | 8.85 ± 0.94 | n = 14 | |
| 5mm (°) | 8.36 ± 0.58 | n = 14 | 4 vs. 5 p=0.79 (post-hoc) |
| **8mm (°)** | **6.80 ± 0.43** | n = 14 | **5 vs. 8 p=0.022 (post-hoc)** |
| 10mm (°) | 6.38 ± 0.37 | n = 14 | 8 vs. 10 p=0.28 (post-hoc) |
| **Tail-on-body mean (°)** |  |  | **p=0.0070, ANOVA** |
| 4mm (°) | 30.24 ± 1.76 | n = 14 | |
| 5mm (°) | 28.86 ± 1.37 | n = 14 | 4 vs. 5 p=0.64 (post-hoc) |
| **8mm (°)** | **24.45 ± 1.76** | n = 14 | **5 vs. 8 p=0.077 (post-hoc)** |
| 10mm (°) | 22.67 ± 1.55 | n = 14 | 8 vs. 10 p=0.19 (post-hoc) |
| Front mean/speed | r=0.300 | n = 14 | p=0.141, linear regression |
| Back mean/speed | r=0.0498 | n = 14 | p=0.715, linear regression |
| **Tail-on-body mean/speed** | **r=0.291** | n = 14 | **p=0.0291, linear regression** |
| Front mean/Centrality | r=-0.147 | n = 14 | p=0.279, linear regression |
| Back mean/Centrality | r=-0.192 | n = 14 | p=0.155, linear regression |
| Tail-on-body mean/Centrality | r=-0.061 | n = 14 | p=0.655, linear regression |

Statistical analysis was performed using ANOVA followed by Bonferroni's post-test. n refers to the number of mice. vs., versus.

### Tail kinematics in narrow-substrate locomotion

Locomoting on any structure narrower than the knees of the mouse is expected to lead to pronounced roll-plane oscillation of the body and possibly a need to compensate for the roll to avoid falling. Thus, we hypothetized that the increasingly lateral position of the tail on thinner ridges (Figure 3E2) would be part of a strategy to provide some counteracting momentum to the hips. To examine this, we tracked the roll-plane motion of mouse tails (Figure 4A1, top) and hips (Figure 4A1, bottom) from the rear camera view while the mice traversed unperturbed across the ridges. Indeed, similarly to the increasingly lateral position of the tail when viewed from the top, the tails (held close to vertical on broadest ridges) descended towards the horizontal alignment when traversing the most challenging ridges while the hips remained on average aligned with horizontal.

**Figure 4.**
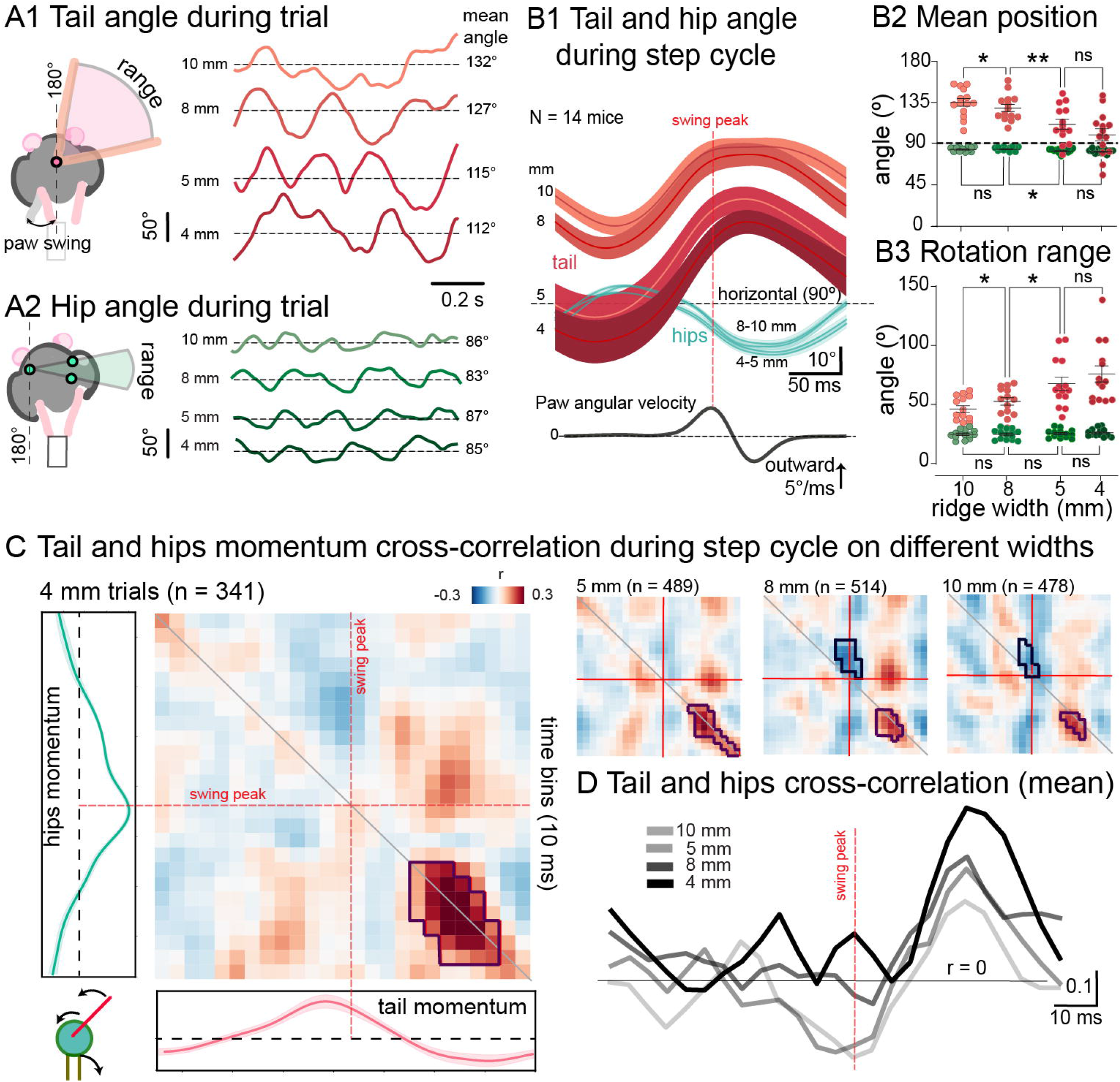
Tail and hip movements during unperturbed ridge traverse. A1 and A2, rear-view angle trajectories of tail (A1) and hip (A2) angles of an example mouse traversing ridges of different widths. One second of the trials is shown starting from the beginning of ridge traverse. Dashed lines indicate the mean position for the tail in a given trial. Schematics on the left depict the measurements. B, Tail and hip movements temporally aligned on the contralateral paw swing. B1, mean ± SEM angle trajectories of tails (red) and hips (green) of 14 mice. Bottom panel shows average of contralateral paw angular velocity, aligned on the outward peak. B2, mean position of the hips and tail. Data are shown as averages of single animals over all trials of a given width. B3, range of tail and hips motion through step cycles. C, cross-correlation between the tail and hip momenta through the step cycles centered on swing peak. Left, cross-correlogram for the 4mm ridge condition. Cross-correlation values are displayed in a normalized intensity range between 0.3 (red) and -0.3 (blue). Cross-correlograms for 5mm, 8mm, and 10mm ridges are shown as smaller panels to the right. “Hotspots” and “coldspots” (see Methods) are depicted with dark contours. Dashed red lines indicate swing peak times. D, Mean correlation values along the diagonal for each ridge width. Darker lines correspond to narrower ridges trials. Red dashed line indicates time of swing peak. * p *<* 0.05, ** p *<* 0.01, ns, p *>* 0.05.

Thus, the tail was repositioned independently from hip alignment, similarly to the lateral displacement discussed before. Furthermore, the oscillatory movement of the tail initial segment increased significantly on narrowing ridges. To investigate whether there is a consistent change in the tail use that could contribute to traversing performance, we examined the oscillatory movements of the tail and hips in temporal alignment with the step cycle (Figure 4B1; see Methods for swing phase definition; see Table 7 for all values and statistical comparisons). Indeed, we found that their oscillation occurs antiphase to each other on all ridges, and both hip and tail angles reach their maximal deviations from horizontal plane during late phase of the contralateral limb swing. This visualization, along with the systematic lowering of the tails (Figure 4B2, red) and increasing amplitude of oscillation (Figure 4B3, red) contrasts the movement of hips that remained remarkably invariant on all ridges (Figure 4B2-3, green).

**Table 7.**
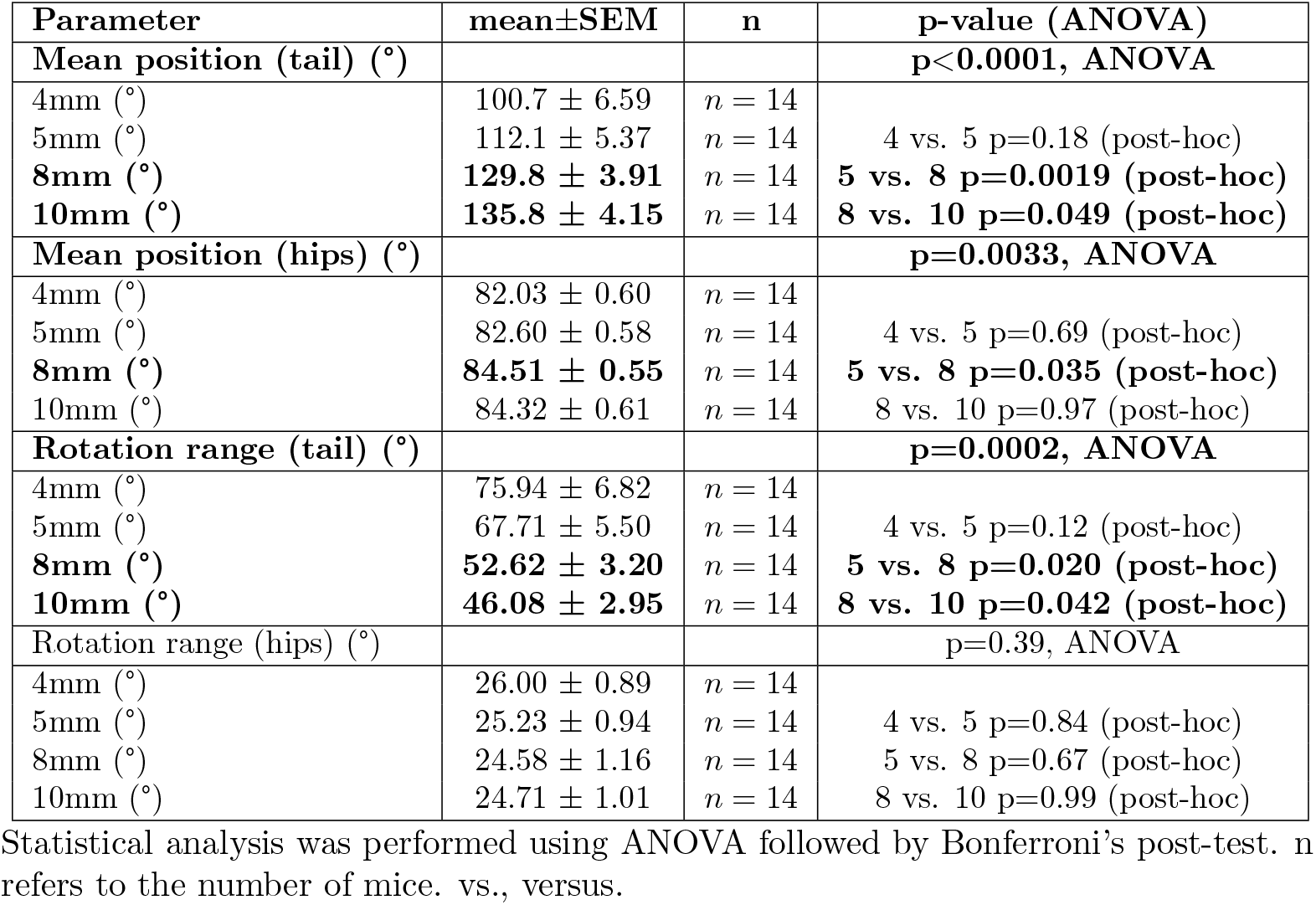
Tail and hips position and rotational range during a step cycle while locomoting on ridges with different width.

Notably, our simple biomechanical model can not provide meaningful estimates for generated angular momenta under the complex and dynamic locomotory context (as opposed to the tilting ridge-context where the magnitude of external perturbation is known). However, it is still insightful to examine the dynamics of tail and hip rotation with respect to each other. Indeed, when examining the cross-correlation between hip and tail oscillations on the 4mm ridge (Figure 4C, left; high correlation region indicated with red pixels and dark outline) we found that their movements are most strongly coupled shortly after the contralateral paw’s stance onset. This coupling is most pronounced when the mouse is traversing the most challenging 4 mm ridge, and diminishes on easier ridges. In contrast, the coupling between the tail and hips develops negative correlation with broader ridges (Figure 4C, right; blue pixels with dark outline), that is centered around the peak swing phase (cross-correlation values shown along the matrix diagonal in Figure 4D). This suggests that mice may flexibly engage their tails to provide appropriate biomechanical support in diverse locomotory contexts.

## Discussion

Here, we presented mice with a naturalistic balancing task: locomotion on a narrow ridge while countering roll-plane perturbations. This setup mirrors the challenges faced by mice in natural environments, such as tree-branch navigation.

Our findings establish that mice initiate high-speed rotational tail movements that deliver substantial angular momentum, largely counteracting roll perturbations (Figure 1). These tail dynamics are only slightly modulated by proprioceptive context defined by the ridge width (Figure 2) suggesting that tail swing, triggered by tilts, might bear resemblance to vestibulo-ocular (VOR) and vestibulo-collic (VCR) reflexes. If confirmed to be vestibularly driven, this tail response could provide an accessible metric — hypothetically termed Vestibulo-Tail Reflex (VTR) — for probing cerebellar and vestibular circuits avoiding the challenges associated with tracking eye or neck movements.

Beyond externally-induced tail movements, we demonstrate purposeful adjustments in hind body and tail posture during unperturbed ridge-crossing. Mice shift their hind-bodies laterally and lower their tails towards the horizontal plane when locomoting on increasingly narrow ridges (Figure 3). Furthermore, the tails engage in phase-locked oscillatory movements with respect to the contralateral hindlimb step cycle (Figure 4). These oscillations and their coupling with hip movements are more pronounced when traversing narrower ridges, even though hip rocking remains largely invariant across different ridge dimensions. It seems possible that the tail is recruited to provide compensatory angular momentum under conditions where the hip/body rotation is not sufficient to ascertain that the center of mass remains safely within the base of support.

There is no doubt that constructing a more detailed model as well as high-resolution tracking of limb and body movements would allow deeper understanding of the forces, torques and context-specificity of the numerous interacting strategies that mice employ to maintain balance. Notably, even without opposable digits in their paws, it is to be expected that mice can generate some amount of stabilizing torque by opposable limb forces depending on the gait used, similarly to primates [15]. However, taking into account the considerable rear-heaviness of mice (COM is located 35.66*±* 1.87 percent closer to the tail base along the body axis), the relative contribution of the front limbs is expected to be rather limited.

### Hierarchy of Balancing Strategies

Maintaining posture and balance is a multilayered endeavor, requiring enactement of context-dependent motor strategies that account for both passive and active self-motion within a dynamic environment [16]. For instance, “change-in-support” strategies (changing position of a limb) often serve as the most efficient mean to restore balance in healthy humans. However, when freedom of foot placement is constrained, “hips-and-ankles” strategies (torque-based shifts of the body’s center of mass) become the primary balancing mechanism [17]. Under the most precarious balancing scenarios such as walking on narrow beams, humans additionally employ upper-body strategies involving arm movements [18–20]. In a sense, we observe here a parallel in that even though mice traversing wide surfaces don’t typically move their tails [7], they do engage their tails under challenging conditions when the freedom of movement is otherwise limited.

Importantly, it must be noted that the mouse tail - like human arms - plays distinct roles under different dynamical circumstances. Experiencing sudden tilt perturbations, the mice stop locomoting and the tail responds to immediate balance threats with swift, high-speed rotations that generate stabilizing angular momentum. On the other hand, during voluntary locomotion where the effective base of support is defined by momentary paw placement arrangement, the tail’s behavior changes markedly. Here, instead of generating large swings, the tail could act as a ‘dynamic rudder’, possibly involved in propelling the animal’s extrapolated center of mass (xCOM [21]) for smooth progression. This suggests a role for anticipatory postural adjustments, wherein the tail’s position and movement are phase-linked to step cycle to assist in upcoming dynamic shifts, possibly influenced by proprioceptive and descending command signals.

Despite these uncertainties we should note that in both scenarios the tail functions as a ‘fifth limb,’ with distinct strategies for reactive balancing and proactive, skillful locomotion. Given these diverse roles, skilled motor coordination in mouse locomotion is incompletely understood without considering the dynamic contributions of the tail. Its versatility in aiding balance and enhancing locomotion underscores its significance in complex motor tasks and potentially provides an accessible avenue for future research into mouse motor control.

### Revising Metrics and Targets for Studying Balance in Rodents

Research on balancing strategies in animals has largely focused on larger species like cats and humans, where lower limbs and trunk movements are the primary focus (but see [22] for the role of tail in cats, and [23] for mentions of tail swings in large primates). A pioneering study by Murray and colleagues [10] has brought attention to balancing strategies in mice, examining hindlimb muscle activity during linearly-accelerating perturbations of the beam mice were traversing. To observe the minuscule movements upon balance perturbations the study was based on EMG recordings from the hindlimb muscles. We propose that the tail could be considered as a novel focus for balancing-related reflex study in mice. Easily visualized and tracked with markerless methods such as DeepLabCut, the tail presents an underexplored yet valuable metric for studying balance in smaller rodents.

Furthermore, we propose using the animal’s estimated Center of Mass (CoM) within its base of support as a direct measure of balance performance. Unlike classic paw-slip metrics ( [8]) that are cumbersome to generate as well as subject to confounding factors such as substrate surface properties, the CoM trajectory provides a nuanced insight into an animal’s balancing strategy and offers a methodologically simple yet robust view of the animal’s balance capabilities [24]. We expect that combining CoM-based balance assessment with other metrics (such as traversing speed and number of stops [25]) may prove to be an additional insightful approach to dissect balancing strategies in the context of specific motor aberrations.

### A short tale of mammalian tails

Even though orienting the body with respect to the gravity vector, maintaining posture as well as differentiating between passive and active self-movement are among the most ancient motor skills of all animals, the transition to terrestrial life and legged locomotion brought about an entirely new dimension to the task of “balancing” - namely, that of maintaining the center of mass (CoM) above the base of support during locomotion. Thus, despite the fact that the vestibular organs and nuclei are among the vertebrates’ most ancient sensorimotor structures that predate emergence of true cerebella in elasmobranchs [26–28], it seems evident that more advanced strategies involving control of the CoM with respect to the instantaneous location of limbs for supporting dynamic erect locomotion (emerging during the Triassic period in both Archosaurs and Therapsids [29]) are necessary.

In this context, it is not surprising that the vestibular nuclei took a significant evolutionary leap between sprawling amphibians and more erect reptiles [30]. Ability to maintain CoM above a narrow base of support would be particularly important in the context of seeking food and refuge among the trees on unstable branches. Notably, the only truly arboreal lizards, chameleons, rely on their firm grip on the branches and slow, large-amplitude limb movements rather than tail positioning to maintain balance on branches narrower than their bodies ( [31]). Thus, the neuromechanical substrates enabling roll-plane stabilizing use of tails (in externally and self-generated contexts) likely evolved uniquely in mammals. This is possibly reflected in the flexible and muscular structure of many mammalian tails, which affords a range of motion and control not seen in the lizards.

It remains unclear why humans lack a tail [32] but the bipedal posture leads to increased demands for controlling CoM sway in the pitch (front-back) rather than roll-plane, diminshing the benefit of a swinging tail. Selection of the postural correction strategies (such as rotational arm movements [33]) involves supraspinal processes in humans similarly to animals [34], and balance performance is potentially linked with interoceptive states, cognitive capacity and age [35–38] reflected in scenarios such as old roadside sobriety check using a balancing task [39]. The intertwined relationship between balance, cognition, and physiology, evident even in such commonplace tests, underscores the importance of delving deeper into these mechanisms.

## Materials and methods

### Animals

All procedures were reviewed and performed in accordance with the OIST Graduate University’s Animal Care and Use Committee (protocol 2020-292-2. C57BL/6 mice were purchased from Clea (Japan) and were acclimatised to the OIST facility for at least 1 week before handling. They were housed in institutional standard cages (5 animals per cage) on a reversed 12-hr light/12-hr dark cycle with ad libitum access to water and food. At the commencement of the handling period, the mice were 8 weeks old.

### Anatomical measurements

In this study, measurements of body and tail mass and length were obtained from six mice carcasses. These mice were of the same sex, strain, and age (10-12 weeks) as those used in the experimental recording. The weight of these mice and the experimental group was compared to ensure the anatomical measurements extracted from the carcasses can be generalized to the experimental group, and no significant difference is shown (t(19)=0.27, p=0.7922). The center of mass (CoM) was determined from these carcasses utilizing the reaction board method, adapting from the method described in [40]. After estmation of the CoM, the tail was dissected at its base from the body and the weight and length of the tail and the body was measured. See Table 8.

**Table 8.**
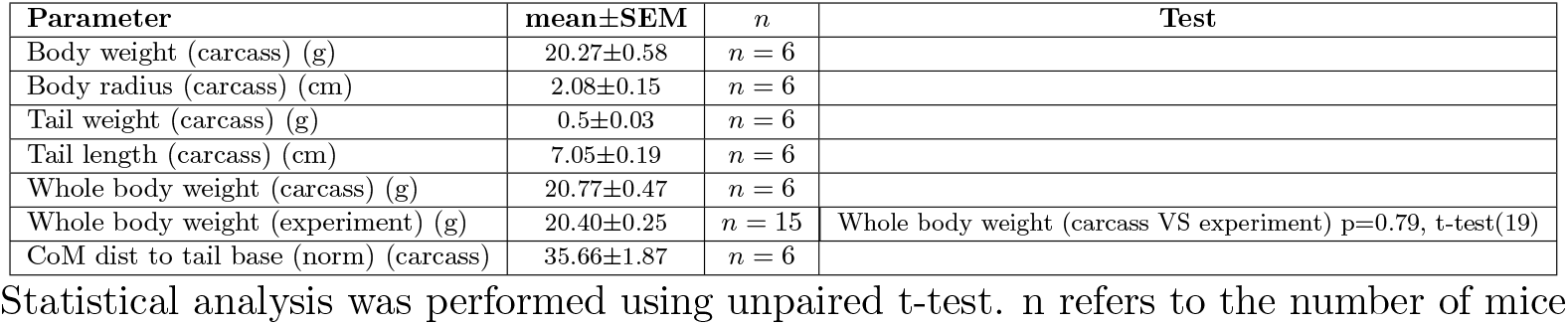
Anatomical measurements.

| Parameter | mean±SEM | n | Test |
| --- | --- | --- | --- |
| Body weight (carcass) (g) | 20.27±0.58 | n = 6 |  |
| Body radius (carcass) (cm) | 2.08±0.15 | n = 6 |  |
| Tail weight (carcass) (g) | 0.5±0.03 | n = 6 |  |
| Tail length (carcass) (cm) | 7.05±0.19 | n = 6 |  |
| Whole body weight (carcass) (g) | 20.77±0.47 | n = 6 |  |
| Whole body weight (experiment) (g) | 20.40±0.25 | n = 15 | Whole body weight (carcass VS experiment) p=0.79, t-test(19) |
| CoM dist to tail base (norm) (carcass) | 35.66±1.87 | n = 6 |  |

Statistical analysis was performed using unpaired t-test. n refers to the number of mice

### The swing task

The mechanical swing comprises a pod designed using Rhynoceros 3D (https://www.rhino3d.com) and 3D-printed using a Ultimaker machine (https://ultimaker.com/3d-printers/). The swing is attached to the shaft of a rotary motor (Teknic CPM-SDSK-2310S-RLN, https://teknic.com) controlled using a custom-written Arduino script. The motor swings the shaft-attached pod so that a back-and-forth motion takes approximately 4 seconds. The mouse is placed in the pod so that it’s tail hangs out. Tail movement was recorded using a high-speed camera (300 Hz, Blackfly S USB3, Teledyne FLIR, Wilsonville, OR, USA). The design files and Arduino scripts for the swing are available at https://github.com/salvatoreNRIM/Ridge_analysis_Nov2023.

### Tilting ridge traverse (TRT) task

The tilting ridge set-up consists of a thin acrylic ridge (4 to 10 mm wide, 50 cm long) attached on one end to a motor (S3003 Servo Motor, Futaba, Japan) controlled by a Bonsai script (https://bonsai-rx.org/) that can tilt the ridge left and right direction with varying amplitudes. A small platform was placed at both ends of the ridge for the mouse to comfortably reside before and after the trial. Mouse movement was recorded at 300 FPS with two high-speed cameras (Blackfly S USB3, Teledyne FLIR, Wilsonville, OR, USA), one placed 50 cm above the ridge and another at the rear end of the ridge. Video recordings were saved in mpg format.

### Animal training

After 2-week handling and habituation period, mice were trained to cross the platform using the 5-mm (days 1, 3, 5 of training) and 8-mm (days 2 and 4) ridges. The mice were gently encouraged to traverse the ridge, using a plastic tube from their home cages placed in front of them as an incentive if necessary. No food or other rewards were used, and the animals were not incentivised artificially (e.g. with food deprivation or stressors). Each training session lasted until the mouse could cross the platform without stops for 10 times in a row. Each training session lasted for approximately 2 hours. Each mouse underwent such training for three consecutive days. On the fourth and fifth day random-direction tilt was introduced in 2 out of 5 crossing trials. All 15 mice used in this study successfully acquired the task and traversed the ridge at the end of the 5th training day.

### Experimental trial structure

During the experiment, mice were subjected to trials with a tilt perturbation randomly intermingled with non-perturbation trials (ratio: 1 perturbation trials per 1 non-perturbation trials), and the onset of the tilt was randomly timed during a trial to prevent anticipatory behavior. After the tilt, the ridge remained in that position through the remaining trial. The baseline perturbation trials involved a 20-degree (190 ms) tilt of the 5-mm ridge in either left or right direction.

Four different ridge widths (4, 5, 8, 10 mm) were used to provide mice with varyingly challenging tasks, as well as a 45-mm, non-tilting ridge that allowed comparison of ridge-traversing locomotion to “flat surface locomotion”. In additional trials the perturbation angle was either decreased or increased (to 10 or 30 degrees (150 or 230 ms), respectively) without changing the angular speed of the tilt. The trials with different ridge widths or tilt amplitudes were randomized according to latin square design [41]. The entire experimental series was completed in 2 weeks (15 experimental days).

### Tracking tail and body movements

Videos were recorded with the top and rear cameras as described above. The nose, tail, hips and hind paw trajectories were extracted using DeepLabCut (version 2.1.7; https://github.com/DeepLabCut/DeepLabCut [42]). A total of 200 image frames (10 videos selected from perturbation/non-perturbation trials, as well as different widths, 20 frames/video) were used to label and train models from both camera views. 90 % of the labeled frames were used for training, and the remaining 10 % for testing. We used a ResNet-50-based neural network for 1,000,000 training iterations, where the cross-entropy loss plateaued to 0.001. We then used a p-cutoff of 0.9 to condition the X,Y coordinates for future analysis. This network was then used to analyze videos.

In addition to the hind body kinematics, position of the body centroid was estimated using the top camera view as the average position of the mouse body silhouette using a custom-made script in Bonsai. Silhouette was extracted by first applying a filter to binarize the image (to separate the region of interest (ROI) from the background), and then extract the ROI centroid, as described in [43] The point in time were the ROI centroid became visible under the top camera were used to extract the time-aligned traces captured by both cameras. Out of this trace the 500 points (centered in time) of the time series were used to compute angles displacement during a trial. Tails and hips angles time-series were either extracted for the trial, or for a step cycle. A step cycle was identified based on the peak of the x projection time-series of the contralteral (with respect to the position of the tail) hind paw marker. This time-point is used to separate the trials based on steps events and project the time-series of hips and tail angles centered in time around this event.

### Kinematics analysis

Custom Python scripts (Python 3.7.4; https://www.python.org run on Windows 10; detailed list of dependencies are listed in the GitHub page https://github.com/salvatoreNRIM/Ridge_analysis_Nov2023) were used to compute angles and instantenous velocities for all DeepLabCut-extracted markers, as well as the speed of the Bonsai-extracted centroid trajectories. All time series were applied a smooth filter (Hanning smoothing) of 10 frames (33 ms) before further processing.

Following parameters were computed from the extracted trajectories:

- **Roll-plane tail angle**: the angle of the initial segment of the tail with respect to vertical as seen from the posterior camera. 180 degrees corresponts to tail pointing up.
- **Yaw-plane tail angle**: the angle of the initial segment of the tail with respect to the ridge, as seen from the top camera. 0 degrees corresponds to tail pointing straight back.
- **Hip angle**: alignment of a line connecting the two hip markers with respect to vertical, as seen from the posterior camera. 90 degrees corresponds to horizontal alignment.
- **Instantenous angular velocity** is the frame by frame gradient of the tail angle.
- **Back angle** is the angle between the tail base, the centroid, and the line parallel to the ridge passing through the tail base. It is used to estimate the mouse back posture while crossing the ridge. 0 degrees corresponds to the hind body being aligned straight back along the ridge.
- **Front angle** is the angle between the centroid, the nose, and the line passing through the nose and parallel to the ridge. 0 degrees corresponds to the head being aligned straight ahead along the ridge.
- **Tail-on-body angle** is obtained by subtracting the back angle from tail angle (determined by tail 2nd marker, tail base and the line parallel to the ridge passing through the tail base). 0 degrees corresponds to the tail being aligned straight back along the ridge.

### Ridge traversing performance assessment

Performance was assessed using 5 metrics: slip count, traversing speed, fraction of trial frames where CoM (Center of Mass) is within Base of Support (BoS), CoM lateral movement and and CoM centrality. Paw slips, defined as the slip of at least one paw from the platform, were manually counted from slow-motion videos acquired at 300Hz. Traversing speed is calculated as the forward distance covered by the centroid (measured from the top camera) divided by the time it took to cover that distance. CoM within BoS for one trial is defined by the fraction of all trial frames where the centroid is within the edges of the ridge. CoM lateral movement during a trial is calculated as the mean absolute distance of the centroid from the midline of the ridge. Finally, CoM centrality is the lateral distance of the centroid normalized by the width of the ridge (where 1 represents the case where the centroid is on the midline and 0 on the edge).

### Biomechanical model

A simple biomechanical model was constructed by approximating the tail to a rod of length 7 cm and mass of 0.5 g, and the body to a cylinder of diameter 2 cm and mass 20 g (weights and size based on weighting 6 male mouse carcasses as described above). The angular momentum for the tail, body, and perturbation were computed with the following formulas:

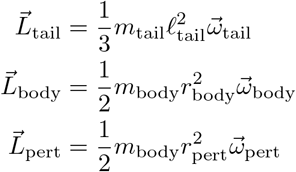

where m is the body mass l is the length of tail.

### Cross-correlation analysis of tail and hip momemta

The cross-correlation was calculated between hips and tail momentum time bins for trials with a certain width. “Hotspots” and “coldspots” were defined by collecting cells with values in positive and negative 5 percentile for the total distribution of correlation and highlighting region with the highest number of contigous cells.

### Statistical analysis

All data are presented as mean ± standard error of mean (SEM). Data were analyzed using one-way analysis of variance (ANOVA), as appropriate. Bonferroni test was used for post-hoc analyses of significant ANOVAs to correct for multiple comparisons. Differences were considered significant at the level of p *<* 0.05. Statistical analysis was performed with Prism 9.0 (GraphPad, San Diego, CA).

## Acknowledgments

The authors would like to thank the OIST Engineering section, all members of the Neuronal Rhythms in Movement team as well as the Transylvanian Experimental Neuroscience Summer School (TENSS) for technical support and discussions. S.A.L. is supported by JSPS DC1 fellowship (202020494). N.I. and M.Y.U. are supported by OIST intramural funding.

## Supporting information

**S1 Video. Tail movement evoked by lateral swinging**. Example of a trial where the mouse is placed in the swing set-up to record tail movements induced by swinging.

**S2 Video. Tail movement evoked by lateral tilting**. Normal and 4x slowed-down videos overlaid with DeepLabCut tracking from tilt trials directed contralaterally and ipsilaterally with respect to the tail position.

**S3 Video. Tail movement evoked by various tilt amplitudes**. Ipsilateral perturbation trials with 10, 20 and 30 degree tilts shown in normal and 4x slowed-down video.

**S4 Video. Tail movement evoked by tilt on ridges of different widths**. Ipsilateral 20-degree perturbation trials using different ridge widths (4, 5, 8 and 10mm) shown in normal and 4x slowed-down video.

**S5 Video. Unperturbed locomotion on ridges of different widths**. Same mouse shown side-by-side on different ridges, in normal and 4x slowed-down video.

